# mRNA destabilisation through CDS-targeting is the primary role of endogenous miRNA in the green alga *Chlamydomonas*

**DOI:** 10.1101/088807

**Authors:** Betty Y-W. Chung, Michael J. Deery, Arnoud J. Groen, Julie Howard, David Baulcombe

## Abstract

MicroRNAs regulate gene expression as part of the RNA-induced silencing complex, where the sequence identity of the miRNA provides the specificity to the target messenger RNA, and the result is target repression. The mode of repression can be through target cleavage, RNA destabilization and/or decreased translational efficiency. Here, we provide a comprehensive global analysis of the evolutionarily distant unicellular green alga *Chlamydomonas reinhardtii* to quantify the effects of miRNA on protein synthesis and RNA abundance. We show that, similar to metazoan systems, miRNAs in *Chlamydomonas* regulate gene-expression primarily by destabilizing mRNAs. However, unlike metazoan miRNA where target site utilization localizes mainly to 3’UTRs, in *Chlamydomonas* utilized target sites lie predominantly within coding regions. These results demonstrate that destabilization of mRNA is the main evolutionarily conserved mode of action for miRNAs, but details of the mechanism diverge between plant and metazoan kingdoms.

## Introduction

MicroRNAs (miRNA) are 21-24 nucleotide RNAs present in many eukaryotes that guide the silencing effector Argonaute (AGO) protein to target mRNAs via a base pairing process (Bartel, 2009). The AGO complex either catalyzes endonucleolytic cleavage or promotes translation repression and/or accelerated decay of this target mRNA (Ameres & Zamore, 2013). There has been controversy about which of these three mechanisms is more significant but recent studies in mammalian cells provide support for accelerated mRNA decay. In ribosome profiling of HEK293 cell-lines transfected with specific miRNAs or of neutrophils with a single miRNA knocked out, Guo et al. demonstrated that miRNA primarily modulates gene expression by destabilizing mRNA instead of repressing translation (Guo *et al*, 2010). Similarly in B and T cells when miR155 is over expressed, the main mechanism for miRNA-mediated gene repression is mRNA destabilization (Eichhorn *et al*, 2014). High-throughput assays with single-cell reporters have also demonstrated that the primary role of miRNA in mammalian cells is to fine-tune gene expression mostly by destabilization of mRNA and mostly through targeting the 3’ untranslated regions (UTR) (Siciliano *et al*, 2013; Schmiedel *et al*, 2015).

In plants there is miRNA-mediated gene regulation (Brodersen & Voinnet, 2009; Reis *et al*, 2015; Li *et al*, 2013) but, unlike metazoan systems, the targets can be in the coding sequence as well as 3’UTR and the mechanism may involve endonucleolytic cleavage rather than accelerated decay or translation inhibition (Brodersen *et al*, 2008; Iwakawa & Tomari, 2013). Most plant studies, however, are based on individual miRNAs or reporter assays and there are few studies in plants on the global effects of miRNA under physiological conditions. We therefore utilized the unicellular green alga *Chlamydomonas reinhardtii*, for which we have previously discovered and characterized its miRNAs (Molnar *et al*, 2007) and generated *DCL3* mutants (Valli *et al*, 2016). As *Chlamydomonas* is evolutionarily divergent from higher plants, miRNA effects observed in both *Chlamydomonas* and, for example, *Arabidopsis* are likely to be general amongst all plants.

*Chlamydomonas* is a particularly amenable experimental system because its unicellularity reduces complications with tissue-specific effects. The *dcl3-1* mutant results in almost complete loss of miRNA as well as 21-nt small interfering (si)RNAs but does not result in obvious growth differences or morphological abnormality under normal conditions (Valli *et al*, 2016). Any effect of *dcl3-1* on gene expression is likely, therefore, to be direct rather than an indirect secondary consequence of metabolic changes due to loss of miRNA-mediated regulation.

Here, through a combination of ribosome profiling, parallel RNA-Seq, sRNA-Seq and quantitative proteomics at mid-log phase of the *dcl3-1* mutant and its corresponding complemented strain we have demonstrated that, in contrast to the metazoan system, the primary effect of miRNA in *Chlamydomonas* is through interaction with CDS regions instead of 3’ UTRs. However, similar to the metazoan system, miRNA in *Chlamydomonas reinhardtii* modulates gene expression primarily by promoting mRNA turnover rather than influencing translation efficiency.

## Results and Discussion

### Loss of DCL3 function does not affect the genome-wide RNA or translation profile

To explore the possibility that DCL3-dependent miRNA or siRNA regulates gene expression by either promoting mRNA turnover or through interfering with translation, we applied ribosome profiling, parallel RNA-Seq and quantitative N15 proteomics to biological triplicates of the vegetative mid-log phase *dcl3-1* mutant and its corresponding complemented derivative (abbreviated as C) carrying a wild type *DCL3* allele introduced into the mutant strain. The experimental protocol is summarized in supplementary Figure 1 and supplementary Figure 2 illustrates the high degree of reproducibility between biological repeats in these data.

**Figure 1.**
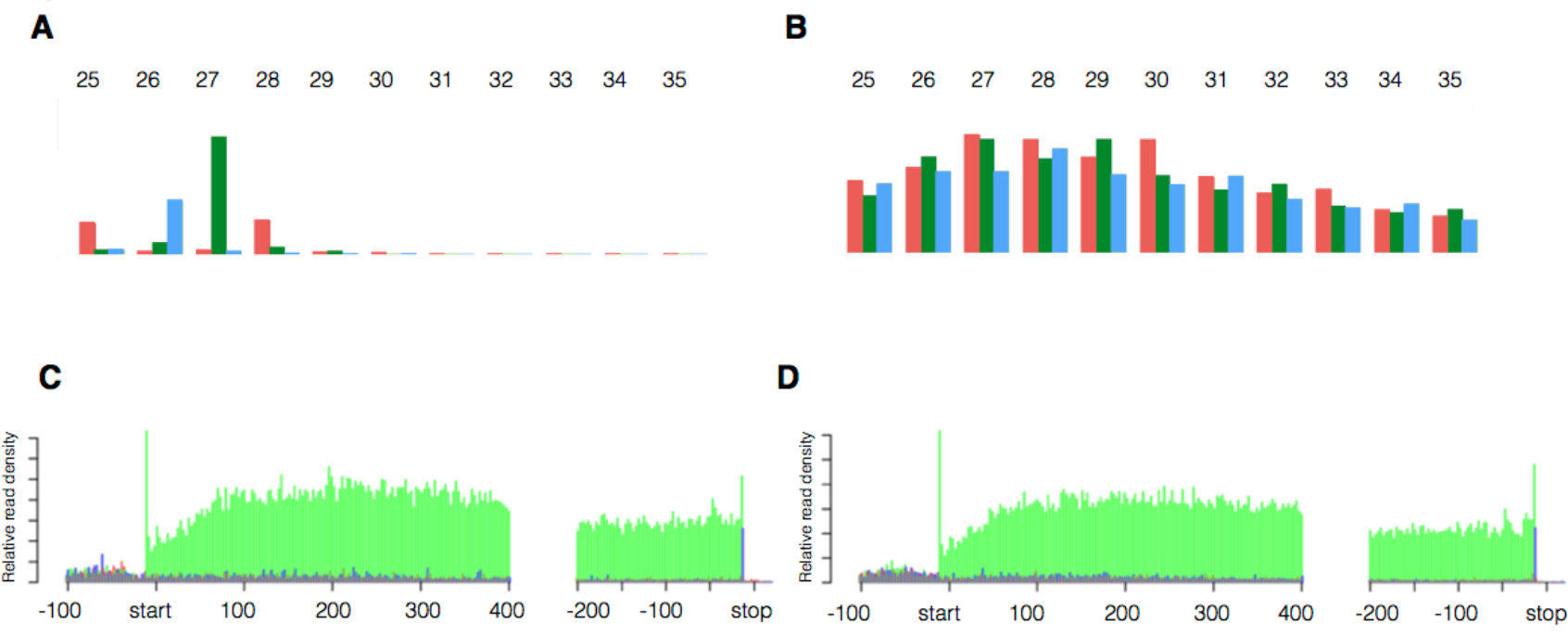
Ribosome profiling data. (A, B) Mapping the 5’ ends of ribosome protected fragments (RPFs) and corresponding RNA-Seq respectively, as a function of read size class (nt), within nucleus-encoded coding ORFs. Red, green and blue bars indicate the proportion of reads that map to codon positions 0, 1 and 2 (respectively). (C, D) 5’ end positions of 27-nt RPFs relative to start and stop codons (nt). Reads were derived from strain C and *dcl3-1* (respectively) and summed over all transcripts. Phasing is indicated using the same colours as in panels A and B.

**Figure 2.**
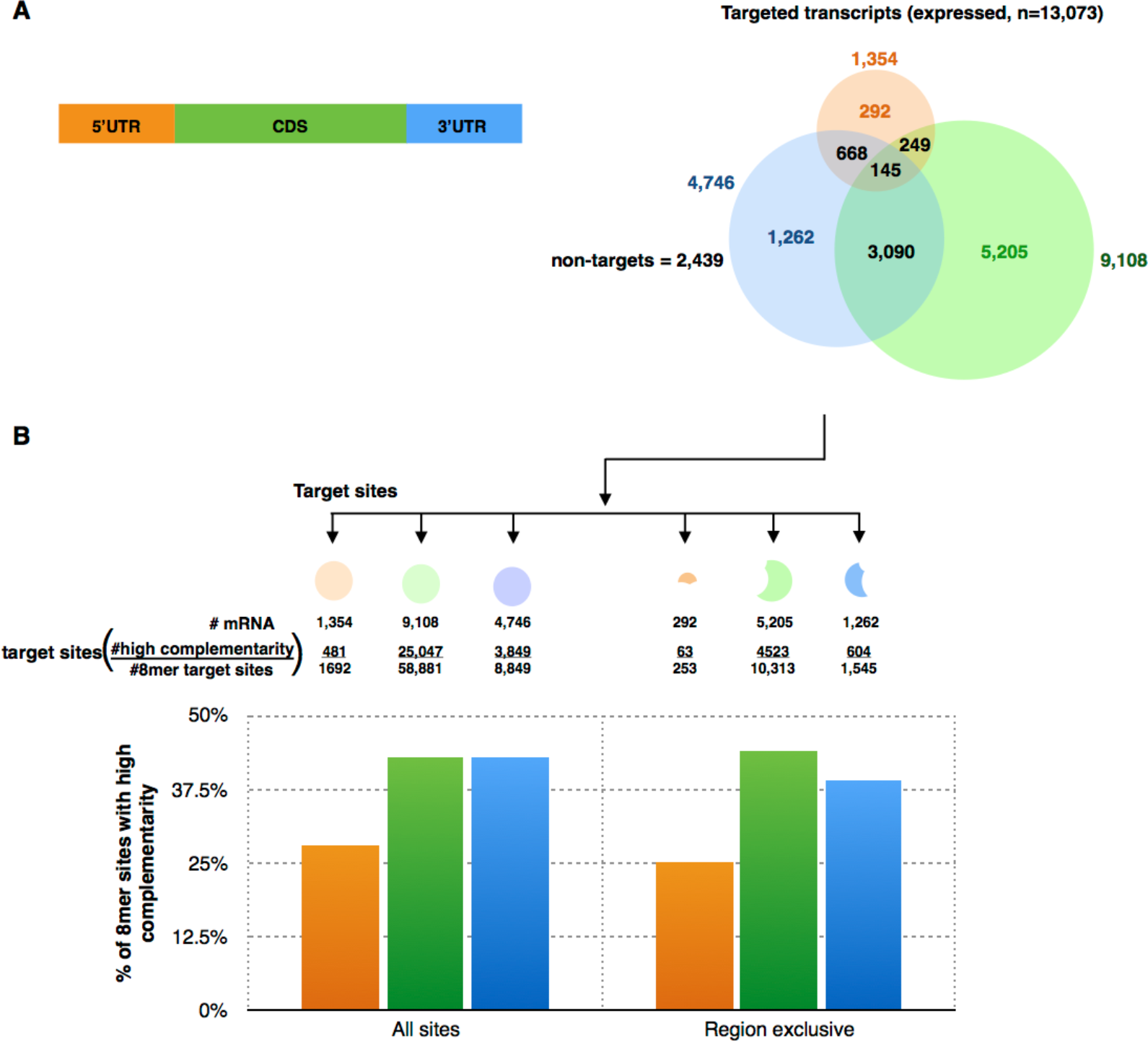
Distribution of 8mer target sites. (A) Venn diagram showing number of transcripts predicted to be targeted with the 8mer rule. (B) Proportion of 8mer target sites that also have at least 50% complementarity from nucleotides 11-21 of the miRNA

The slightly smaller footprint size of plant/algae ribosomes leads to differences in the phasing patterns compared to mammalian ribosome profiling studies (Chung et al., 2015). In both the complemented strain C and the *dcl3* mutant, the 5’ end of the 27-nt ribosome protected fragments (RPFs), mapped predominantly to the second codon position; in contrast, and as expected, RNA-Seq reads were uniformly distributed at all three codon positions (Figures 1A and B). The RPF 5’ end position distributions at start and stop codons were also similar in the *dcl3-1* and C strains (Figures 1C and D respectively) in that there was a sharp 27-nt peak on the start codon (reflecting the rate-limiting initiation step of translation) and a sharp 28-nt peak on the stop codon (reflecting the conformation change from an elongating ribosome to a terminating ribosome) (Chung *et al*, 2015), and the expression level of DCL3 are similar within the triplicates of either the complement or *dcl3-1* mutant background (Supplementary Figures 3A-C). Ribosome protected fragments (RPF), RNA abundance (RA), and translational efficiencies (TE) for expressed genes are well correlated between *dcl3-1* and C (*R*^2^ = 0.95, 0.97 and 0.98 for TE, RPF and RNA, respectively, Supplementary Figure 3D). From these data, we conclude that any global effect of DCL3 on the translatome is minor. Nevertheless, this analysis involved all mRNAs and any quantitative effects on the subset of RNAs with miRNA target motifs may have been masked.

**Figure 3.**
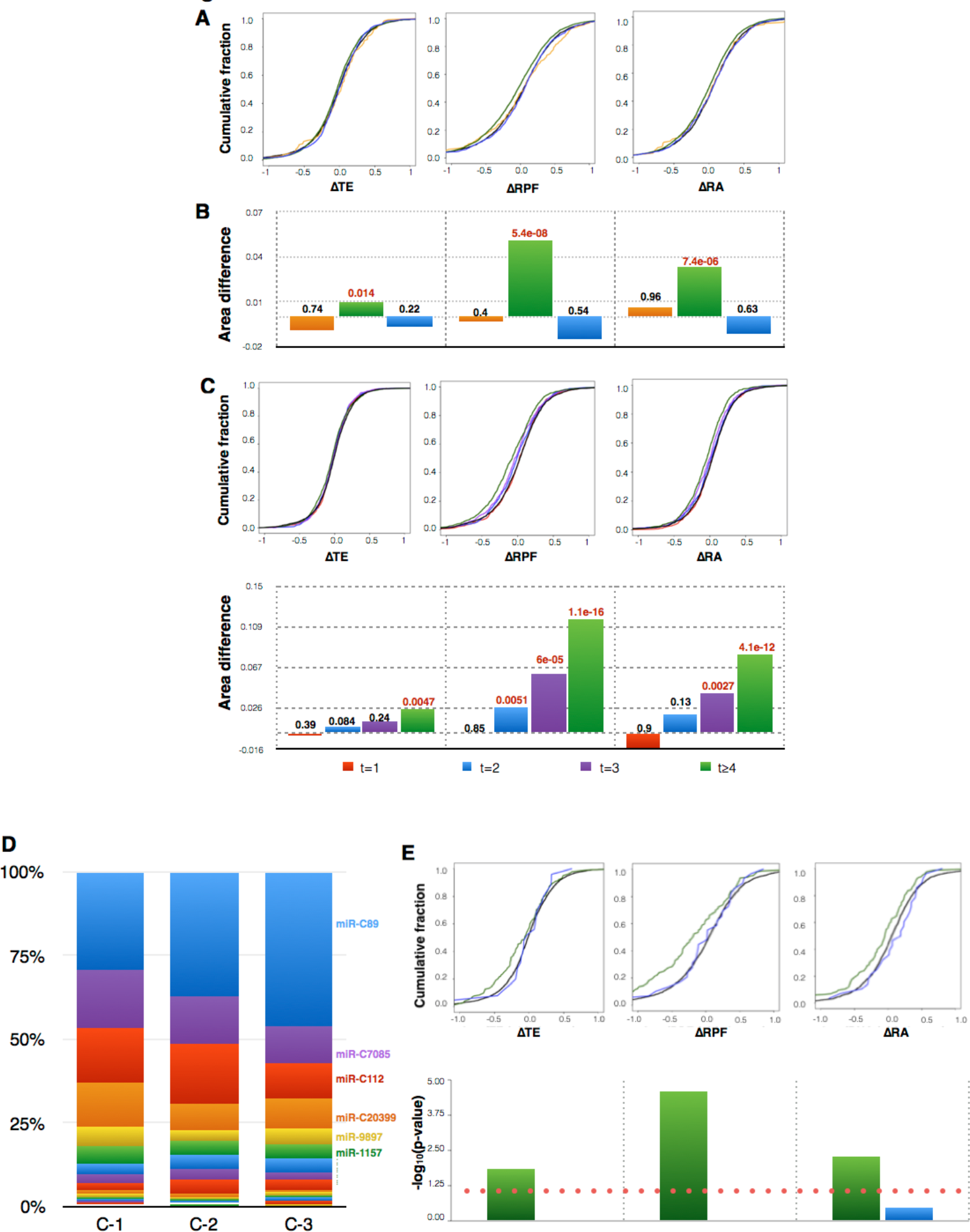
miRNA downregulates gene expression primarily through mRNA destabilization by CDS targeting. (A) Cumulative distributions of ΔTE (left), ΔRPF (middle) and ΔRA (right) log_2_ fold changes in *dcl3-1* relative to C. Colours correspond to genes containing predicted 8mer miRNA target sites exclusively in the 5’UTR (orange), CDS (green), 3’UTR (blue), or no targets (black). (B) Bar graph of differences between area under cumulative distribution of mRNA containing target sites and non-target containing mRNAs (5’UTR, CDS and 3’UTR in orange, green and blue, respectively). Significance (K.S. test) of the area differences are indicated above each bar; p-values less than 0.01 are highlighted in red. (C) Cumulative *dcl3-1* relative to C log_2_ fold change distributions of ΔTE, ΔRPF and ΔRAA in mRNAs with 0 (black), 1 (red), 2 (blue), 3 (purple) or 4 or more (green) CDS-exclusive target sites. (D) Bar graph of differences between area under cumulative distribution of mRNA containing 1 (red), 2 (blue), 3 (purple) or 4 or more (green) CDS-exclusive target sites and non-target containing mRNAs. Significance (K.S. test) of the area differences are indicated above each bar; p-values less than 0.01 are highlighted in red. (E) Normalised miRNA abundances in three biological replicates. (F) Cumulative distributions (top) and significance (bottom) of ΔTE (left), ΔRPF (middle) and ΔRA (right) log_2_ fold changes for mRNAs containing miR-C89 target sites exclusively within the CDS (green) or 3’UTR (blue) (sample sizes 141 and 25, respectively).

To explore this possibility, we refined our analysis by dividing the mRNA profiles into those with or without predicted targets of the DCL3-dependent miRNAs. The first stage in this analysis was to re-evaluate the miRNA precursors in *C. reinhardtii* that we had previously identified as being both coding and non-coding RNAs. Now, however, with the use of the RPF data to identify translated open reading frames, we find that all miRNAs in this alga derive from introns or the exons (3’UTR or coding) of mRNAs. Supplementary table 2 is an updated summary of the 42 miRNA precursors in *C. reinhardtii* described in Valli et al (2016).

Our subsequent analysis differentiated mRNAs with miRNA targets in the 5’ UTR, CDS and 3’ UTR from those without targets. The CDS regions were defined by the R software Bioconductor package – riboSeqR - that utilizes the triplet periodicity of ribosome profiling for the *de novo* inference of AUG-initiated coding sequences that are supported by RPFs (Chung *et al*, 2015) and we used the seed-sequence rule to identify miRNA target motifs (Lewis *et al*, 2003; Agarwal *et al*, 2015). This rule requires base-pairing of the first 8 nucleotides of miRNA and it is supported by direct assay of miRNA targeting and structural studies of human AGO2 (Schirle *et al*, 2014) and by experimental tests in higher plants (Mallory *et al*, 2004) and *C. reinhardtii* (Yamasaki *et al*, 2013).

To identify the miRNA-target mRNAs we first looked for the most abundant miRNAs based on our sRNA-Seq data and filtered for the 19 most-abundant *DCL3-*dependent miRNAs (Supplementary Figure 5; see also Materials and Methods). Using these, we then applied the TargetScan prediction algorithm (Lewis *et al*, 2003; Agarwal *et al*, 2015) to the mRNAs with RPF-validated ORFs. This criterion meant that the TargetScan algorithm was applied to 13,073 expressed transcripts (out of 17,741 annotated transcripts) of which 2,439 do not contain any predicted 8mer miRNA target sites. Of all the predicted target sites, a larger proportion (70%) are located in the CDS (Figure 2A) compared to UTRs (10% for 5’UTR and 36% for 3’UTR). This distribution is likely, at least in part, a reflection of greater length of the CDS compared to UTR regions. Using a more stringent miRNA targeting rule did not have a large change on these numbers: a significant portion of the mRNAs with seed sequence targets also have >50% sequence complementarity to the target mRNA in the sequences downstream of the 5’ eight nucleotides (Figure 2B).

Next, we excluded the RNAs with predicted target sites in more than one region (5’UTR/CDS/3’UTR) because for these it would have not been possible to differentiate the effects of miRNA acting in the different regions. In addition, we also excluded mRNAs with miRNA precursors because they are unstable in the presence of DCL3 as a consequence of miRNA processing (see supplementary Figure 6 and (Valli *et al*, 2016)). Following application of these filters our further analysis was based on 292 mRNAs with 5’ UTR targets, 5,205 with CDS targets, 1,262 with targets in the 3’ UTR and the 2,439 without predicted targets.

To assess the miRNA-mediated effects of DCL3 we plotted cumulative distributions of differential translation efficiency, total RPF and RA for target and non-target mRNAs in the *dcl3-1* mutant and C (Figure 3A). Differential TE is computed as (RPF_C_/RNA_C_)/(RPF*_dcl3_*/RNA*_dcl3_*). The analysis revealed that, similar to the analysis of mammalian cells and zebrafish (Guo *et al*, 2010; Bazzini *et al*, 2012), the major effects of Dicer loss of function (*dcl3-1* vs C) were in the RPF and RNA data but not in TE. The effects were evident as a shift to increased RNA abundance for mRNAs with target sites in *dcl3-1* and they are consistent with the canonical role of miRNAs as negative regulators.

The difference in *dcl3-1* versus C was greater in transcripts with CDS rather than UTR target sites and it was dependent on the presence of miRNA target sequences (Figure 3A and B). The mRNAs with four or more CDS targets were affected to a greater extent than those with fewer target sites (Figures 3C). Furthermore, these effects are also consistent at the protein level for mRNAs with supportive proteomics data (Supplementary Figure 7). The global effect of mRNA repression is not likely due to cleavage as there are only 85 potential CDS-target sites (83 mRNAs) complying with the plant targeting rule that is utilized by *Chlamydomonas* for target cleavage (Molnar *et al*, 2007). Moreover, of these potential cleavage site targets within CDS, only 18/83 mRNAs were expressed in our dataset. There was no significant differential effect on TE or RA between *dcl3-1* and C for these mRNAs. A recent degradome study is also consistent with there being minimal miRNA target site cleavage in *Chlamydomonas*. The study involved miR-910, an miRNA also expressed in our sample, that cleaved only two mRNAs upon salt-stress (Gao *et al*, 2016). The endogenous miRNA-mediated RNA down-regulation by CDS-targeted miRNA is not, therefore, likely to be mainly through target cleavage.

Finally, we tested the effect of miRNA abundance on TE, RPF and RA by focusing on the most abundant miRNA in our corresponding sRNA-Seq datasets: miR-C89 (Figure 3D, E and supplementary Figure 5; 5’UTR and protein data excluded due to small sample size). MiR-C89 correlated with a larger shift in TE and RA than other miRNAs consistent with magnitude of the effect being influenced by miRNA abundance.

From these findings we conclude that, similar to metazoan systems (Guo *et al*, 2010; Eichhorn *et al*, 2014), *Chlamydomonas* miRNA generally fine tunes gene expression through an effect on RNA abundance rather than translation efficiency (Figure 3). The global effect was small (Figures 3A and B), as in metazoans (Guo *et al*, 2010). Unlike metazoans, however, the primary targets of miRNAs in *Chlamydomonas* are in the CDS instead of 3’UTRs (Figure 3). This difference may reflect differences between *Chlamydomonas* and metazoans in the ways in which miRNAs may influence elongating ribosomes.

### Translation efficiency of 80S ribosomal proteins is higher in the DCL3 mutant

Our finding that miRNA targeting in *Chlamydomonas* is influenced by miRNA abundance and the number of target sites (Figure 3) implies that some mRNAs may be affected more than others. Therefore, to detect possible changes in individual mRNAs we plotted the *dcl3-1* versus C differences in TE and RA for all mRNAs with CDS-exclusive targets sites (Figure 4). Using DCL3 as a benchmark (log2FC(TE)=0.7 and log2FC(RNA) = 1.18), individual RNAs that are negatively regulated by miRNAs would distribute in field A of this figure if TE is affected (i.e. log2FC(TE) ≤ -0.7, yellow shaded area), field C if RA is affected but not TE (i.e. log2FC (RA) ≤ -1.18, -0.7 ≤ log2FC(TE) ≤ 0.7, purple shaded area) and in field B if there was a double effect on both TE and RA (log2FC(RA) ≤ -1.18, log2FC(TE) ≤ - 0.7, red shaded area). Corresponding positive regulation would be indicated by distribution in fields A’, B’ and C’ respectively (Figure 4A).

**Figure 4.**
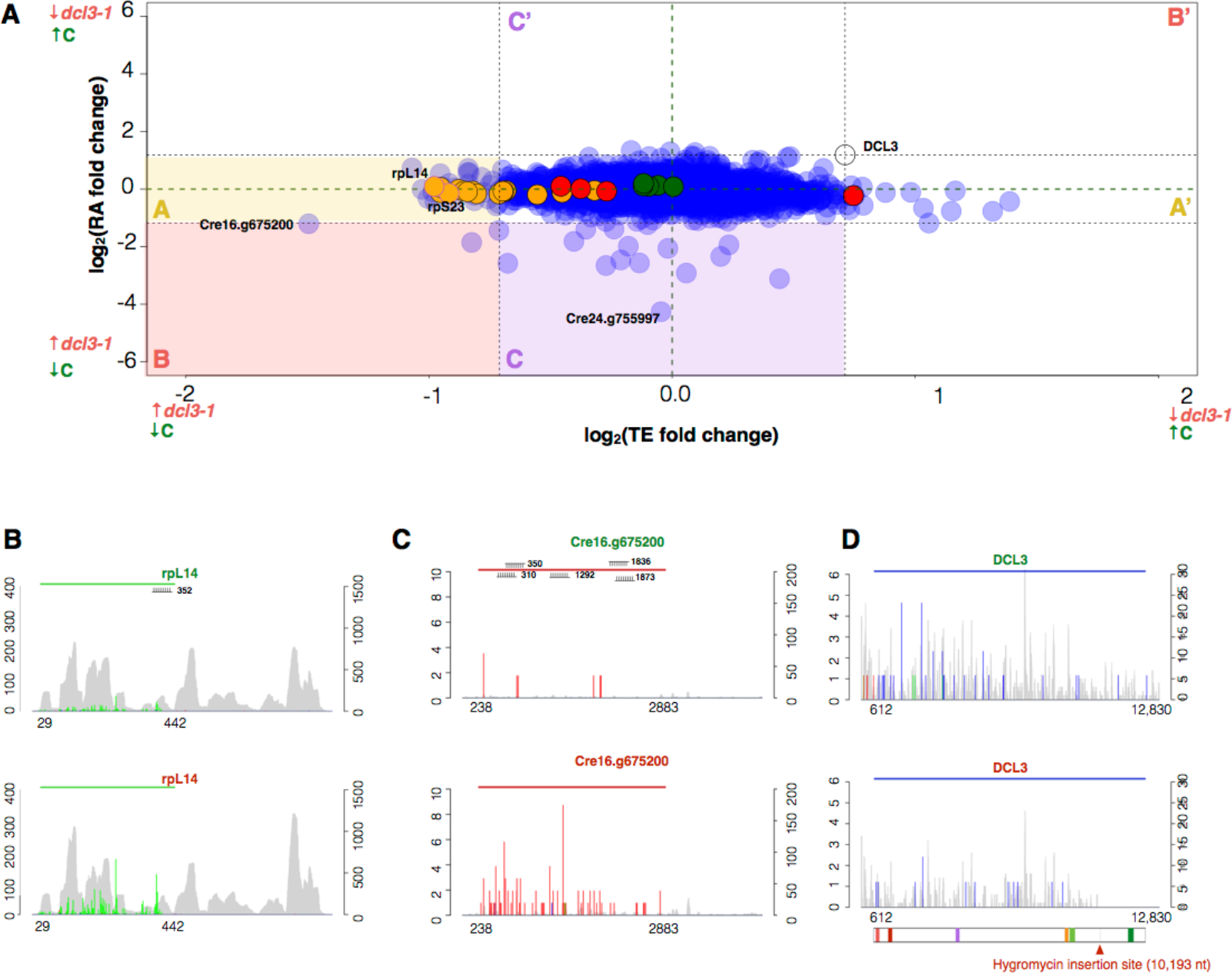
Effects of miRNAs on TE and RA. (A) Correspondence between TE and RA fold-changes between *dcl3-1* and C for nuclear-encoded genes containing miRNA target sites exclusively within the CDS (except DCL3, which was included as a marker). 80S, chloroplast and mitochondria ribosomal proteins are in orange, green and red, respectively. (B-C) Histograms of 5’ end positions of normalised RPF (coloured, left-axis) and RNA-Seq (grey, right-axis) 27-nt reads mapped to genes with high differential TE: ribosomal proteins rpL14 and Cre16.g675200. The top (green title) and bottom (red title) graphs are derived from either the complement or *dcl3-*1 allele, respectively. The coloured horizontal line indicates the riboSeqR *de novo*-defined ORF; positions of potential miRNA target sites are annotated. (D) Histogram of 5’ end positions of normalized RPF (coloured, left-axis) and RNA-Seq (grey, right-axis) 27-nt reads mapped to DCL3 transcripts. The top and bottom graphs are derived from either the complement or *dcl3-1* allele, respectively. The blue horizontal line indicates the re-annotated CDS (612-12,830 nt). The schematic below the plot shows the domain organization of DCL3 which contains two DEAD/DEAH box helicase domains (light and dark red boxes), a helicase C domain (purple box), a proline-rich domain (orange box) and two ribonuclease III domains a and b (light and dark green boxes, respectively). The thick grey line and the corresponding red arrow indicate the hygromycin insertion site (nt 10,193).

The distribution of mRNA in this plot is consistent with a higher degree of negative rather than positive regulation on a few mRNAs: there were 32 and 16 targets in A and A’ respectively, 3 and 0 in B and B’ and 15 and 3 in C and C’. From this analysis we conclude that there may be up to 32 mRNAs that are subject to translational regulation by miRNAs (from the A and B fields), 15 subject to regulation of RNA abundance (from the B and C fields) and 3 subject to regulation at both levels. The RNA-Seq and RPF data for *DCL3* mRNA and selected miRNA targets including the 3 from field B are presented in Figure 4 B-G.

It is striking that mRNAs subject to either translational or RNA stability regulation (i.e. field A and C) are enriched with those encoding RNA-interacting proteins (e.g. translation, transcription and rRNA processing) (Supplementary Table 3). Of the mRNAs subject to translational regulation a gene ontology analysis revealed the enriched pathway of “translation and ribosome” with the mRNAs for 80S ribosomal proteins being particularly prominent (Figure 4 and Supplementary Table 3). These candidates also contribute to the outlier group for TE and RPF but not RA in the cumulative distributions for transcripts with supporting proteomic data (Supplementary Figure 7). However, we do not observe enrichment for this pathway in previously published mammalian datasets (Guo *et al*, 2010) of miR-233 knockout cultured neutrophils compared with wild-type culture neutrophils, and HeLa cells after transfection with miR-1 or miR-155 (Supplementary Figure 8).

The enrichment of “translation and ribosome” function in fields A and C of Figure 4A is specific for 80S ribosomal proteins; the nucleus-encoded 70S ribosomal proteins for both chloroplasts and mitochondria were an internal control and cluster around the 0-fold change axis for both TE and RNA (Figure 4A). It is likely therefore that the specific effect for the 80S factors reflects the targeting specificity of miRNAs in *Chlamydomonas* or that it is a compensatory mechanism for the loss of a layer of regulation in the *dcl3-1* mutant.

It is possible that the distribution of ribosomes on the mRNA would be affected by absence of miRNAs (see Figures 4B and C for example rpL14 and Cre16.g675200.t1). However, we did not observe any significant correlation between the position of the miRNA target sites and the distribution of RPF or RNA reads for the mRNAs of fields A and C of Figure 4A either individually or through a global analysis of multiple RNAs. In contrast, in the mRNA for *DCL3* there was an effect: the RPFs in the C sample extended to the stop codon and the RNA-Seq reads covered the full length mRNA whereas, in *dcl3-1*, the RPF and RNA-Seq data were more sparse than in C and they stopped at the site of the mutagenic *hyg* insert (Figure 4D and Supplementary 3C). Clearly, from this *DCL3* analysis, the RPF and RNA-Seq data can reflect both the quantitative and qualitative aspects of ribosome distribution and RNA accumulation.

That there was no significant change of RPF surrounding miRNA target sites indicates that RISC does not induce ribosome pileup in CDS regions. Presumably the efficient RNA helicase activity of the ribosomes is able to overcome the steric hindrance by the RISC in *Chlamydomonas* (Korostelev *et al*, 2006; Qu *et al*, 2011). There may, however, be a transient effect on ribosome translocation. Having now identified these RNAs with the greatest effect on TE and RNA we will be able to explore the factors affecting the two modes of RNA regulation and the conditions under which miRNAs have the greatest effect on their RNA targets.

## Materials and Methods

Three independent fresh single colonies of *Chlamydomonas reinhardtii* cells were sub-cultured as biological triplicates. Cells where grown in 50 ml Tris-acetate-phosphate (TAP) medium at 23 ºC in baffled flasks on a rotatory shaker (140 rpm) under constant illumination with white light (70 µE m^2^ sec^-1^) to mid-log phase (OD_750_ ∼ 0.6), followed by inoculation into 750 ml TAP in 2 L baffled flasks at OD_750_ = 0.2. These were cultured in the same conditions until mid-log phase prior to harvesting by filtering off the media, after which the cell paste was immediately flash frozen and pulverized in liquid nitrogen with 5 mL of pre-frozen buffer (20 mM Tris-Cl pH 7.5, 140 mM KCl, 5 mM MgCl_2_, 10 µg/ml cycloheximide, 100 µg/mL chloramphenicol, 0.05 mM DTT, 0.5% NP40, 1% Triton X-100 and 5% sucrose). The frozen powder was gradually thawed on ice and clarified by centrifugation for 30 min at 4700 rpm at 4 ºC followed by adjustment of A_254_ = 10 before further treatment, or snap frozen in liquid nitrogen and stored at −80 ºC.

### Metabolic labelling and LC-MS/MS

For metabolic labelling, ammonia chloride (14N) was replaced with ammonia chloride-15N (Cambridge Isotope Laboratories Inc) in the TAP media used to maintain *dcl3-1*. There were no obvious differences in growth rates between algae maintained in N14 and N15. *dcl3-1-*N15 and *Complement-*N14 were mixed equally prior to protein extraction via TCA-acetone precipitation followed by resuspension in resuspension buffer (8 M urea, 500 mM NaCl, 10 mM Tris-Cl pH 8, 5 mM DTT) and resolved in 1.5 mm 10% bis-tris Novex Gel (Thermo Fisher Scientific Inc, Waltham, MA, USA). The experiment was performed in biological triplicate.

1D gel bands (12 per lane) were transferred into a 96-well PCR plate. The bands were cut into 1 mm^2^ pieces, de-stained, reduced (DTT), alkylated (iodoacetamide) and subjected to enzymatic digestion with trypsin overnight at 37 °C. After digestion, the supernatant was pipetted into a sample vial and loaded onto an autosampler for automated LC-MS/MS analysis.

All LC-MS/MS experiments were performed using a Dionex Ultimate 3000 RSLC nanoUPLC (Thermo Fisher Scientific Inc, Waltham, MA, USA) system and a QExactive Orbitrap mass spectrometer (Thermo Fisher Scientific Inc, Waltham, MA, USA). Separation of peptides was performed by reverse-phase chromatography at a flow rate of 300 nL/min and a Thermo Scientific reverse-phase nano Easy-spray column (Thermo Scientific PepMap C18, 2 µm particle size, 100 Å pore size, 75 µm i.d. x 50 cm length). Peptides were loaded onto a pre-column (Thermo Scientific PepMap 100 C18, 5 µm particle size, 100 Å pore size, 300 µm i.d. x 5 mm length) from the Ultimate 3000 autosampler with 0.1% formic acid for 3 min at a flow rate of 10 µL/min. After this period, the column valve was switched to allow elution of peptides from the pre-column onto the analytical column. Solvent A was water + 0.1% formic acid and solvent B was 80% acetonitrile, 20% water + 0.1% formic acid. The linear gradient employed was 2-40% B in 30 min (total run time including a high organic wash step and requilibration was 60 min).

The LC eluant was sprayed into the mass spectrometer by means of an Easy-Spray source (Thermo Fisher Scientific Inc.). All *m/z* values of eluting ions were measured in an Orbitrap mass analyzer, set at a resolution of 70000 and was scanned between *m/z* 380-1500. Data dependent scans (Top 20) were employed to automatically isolate and generate fragment ions by higher energy collisional dissociation (HCD, NCE:25%) in the HCD collision cell and measurement of the resulting fragment ions was performed in the Orbitrap analyser, set at a resolution of 17500. Singly charged ions and ions with unassigned charge states were excluded from being selected for MS/MS and a dynamic exclusion window of 20 s was employed.

### Protein identification and relative quantitation

Data were recorded using Xcalibur™ software version 2.1 (Thermo Fisher Scientific, San Jose, CA). Files were converted from .raw to .mzXML using MSConvert and then .mzXML files to .mgf using the in-house software iSPY (Gutteridge *et al*, 2010; Marondedze *et al*, 2016). The .mgf files were submitted to the Mascot search algorithm. The following parameters were employed: carbamidomethyl as a fixed modification, and oxidation on methionine (M) residues and phosphorylation on serine (S), threonine (T), and tyrosine (Y) residues as variable modifications; 20 ppm for peptide tolerance, 0.1 Da of MS/MS tolerance; a maximum of two missed cleavages, a peptide charges of +2, +3, or +4; and selection of a decoy database. Mascot .dat output files were imported into iSPY for 14N/15N quantitation and analysed through Percolator for improved identification (Brosch *et al*, 2009). The 14N and 15N peptide isotopic peaks from the MS1 dataset were used to compare the theoretical mass difference between the heavy and light peptides, and the typical isotopic distribution patterns. Only unique peptides with a posterior error probability (PEP-value) of ≤ 0.05 were considered for further analysis. Spectra were merged into peptides and proteins based on their median intensity in MS1, meaning the more intense the signal of the spectrum, the more weight it added to quantitation. The statistical programming environment R was used to process iSPY output files to check for the 15N incorporation rate and to confirm that the data were normally distributed. After normalization, only peptides detected in at least two biological replicates, with a fold change ≥ 1.5 and a *p*-value ≤ 0.05 were considered for further analysis. Relative protein expression values were computed as (Protein_C_/Protein_dcl3_) using the average of the triplicates for all follow-up analysis.

### Nuclease footprinting

Lysates (200 µL) were slowly thawed on ice and treated with 6000 units RNase I (Thermo Fisher Scientific Inc.). in a thermo-mixer at 28 ºC, 400 rpm for 30 min. The reaction was stopped by mixing the digest reaction with 120 units of SUPERase-In RNase inhibitor (Thermo Fisher Scientific Inc.) followed by centrifugation for 2 min at 14000 rpm at 4 ºC to further clarify any remaining debris. The supernatant was layered onto a 1 M sucrose cushion prepared in *Chlamydomonas* polysome buffer, and RNA were purified as described in Ingolia et al (Ingolia *et al*, 2009).

### Ribosome profiling and RNA-Seq

The methodologies were largely based on the protocols of Ingolia et al and Guo et al (Ingolia *et al*, 2009; Guo *et al*, 2010) with modifications (i) mRNA for corresponding RNA-Seq was enriched by removal of rRNA using the ribo-zero kit (plant seed and root kit), (ii) RNA-Seq size selection was in parallel with ribosome profiling (i.e. between 26 and 34 nt), and (iii) for ribosome profiling, ribosomal RNA contamination was removed by two rounds of treatment with duplex specific nuclease (DSN) for 30 min as described in Chung et al (2015).

### Preparation for sRNA libraries

Small RNA from total RNA samples used for RNA-Seq were size excluded in 15% TBU gel for miRNA enrichment (Thermos Scientific). The sRNA were further prepared according to the NEXTflex small RNA-Seq kit v2 (Bio Scientific), followed by sequencing on the NextSeq500 platform.

### Computational analysis of ribosome profiling and RNA-Seq data

After removal of adaptor sequences, Illumina sequencing reads were mapped to the reference transcriptome (Phytozome 281) or miRNA precursor sequences described in Valli *et. al*. 2016 using bowtie-1 and processed as described in Chung *et. al*. 2015. Only mRNAs with more than 50 RPF reads of size 27 or 28 nt uniquely mapped to more than 10 positions were considered. Corresponding RNA-Seq reads within coding regions *de novo* defined by ribosome profiling were extracted for differential RA as well as TE analysis using riboSeqR as described in Chung *et. al*. 2015. Further filtering was applied for fold change analyses where mRNAs were only considered if they had (i) at least 10 normalised RPF and 10 normalised RNA counts, and (ii) the sum of all RPF or RNA counts over the three biological replicates for both *dcl3-1* and complement combined is at least 200. Normalisation was based on BaysSeq output (Hardcastle & Kelly, 2010). Cumulative distributions for TE, RPF and RA fold changes were calculated based on the average of all three replicates. Differential analyses for the mouse data were obtained from the Gene expression Omnibus in NCBI (Guo *et al*, 2010) (accession:GSE220001 and GSE21992).

### Target prediction

Target prediction was done using TargetScan (Agarwal *et al*, 2015) using the same transcriptome input as for the ribosome profiling analysis. As there are no conserved sites available due to lack of miRNA data from the green algae phylum, we could not calculate context and scores; thus we only utilized the part of the software to detect all possible miRNA target sites. Further, as the efficacy between 8mer-A1 and 8mer-m8 sites are similar, we combined both types of target sites in the 8mer prediction, similar to Guo et al. (2010) and Agarwal et al. (2015). Target prediction based on the plant rule was performed via TAPIR (Bonnet *et al*, 2010).

The list of miRNA used was based on the 19 *DCL3-*dependent miRNAs expressed based on the sRNA data, where the average reads within the complement is greater than 400 and the average ratio of complement to *dcl3-1* reads is greater than 150. The selected *DCL3-*dependent miRNA used are: chromosome_5_3227666_3227753_+ (miR-C89), chromosome_6_6776108_6776193_+ (miR-cluster20399), chromosome_13_2001067_2001197_- (miR-cluster 7085), chromosome_10_3399870_3399999_- (miR9897), chromosome_13_3152367_3152452_- (miR-C112), chromosome_6_3067368_3067456_+ (miR1162), chromosome_12_6402226_6402307_-(miR1157), chromosome_9_6365928_6366014_- (miR912), chromosome_7_4386252_4386309_-, chromosome_17_6144120_6144204_+ (miR-cluster12551), chromosome_1_7070552_7070605_-, chromosome_16_185088_185174_-(miR1169), chromosome_2_8349161_8349264_+, chromosome_2_9129508_9129593_-miR-cluster14712), chromosome_7_5926395_5926482_+ (miR-C59), chromosome_14_3218783_3218866_- (miR910), chromosome_6_7063792_7063881_- (miR1152), chromosome_4_3100624_3100751_+ (miR1153) and chromosome_1_5106349_5106475_+ (miR-C82). The miRNA precursor sequence used for mapping was based on Valli et al (2016). Only 8mer sites were utilized, and 8mer complementarity was verified via extraction of target sites followed by miRNA complementarity assessment using the Vienna RNA package program RNAduplex. The level of 3’ complementarity was similarly investigated where nt 9 to 21 of the target site 3’ of the seed region was extracted and the level of complementarity assessed with RNAduplex.

## Acknowledgements

We thank J. Barlow for technical assistance and media preparation; B. Santos for technical bioinformatic support; and T. J. Hardcastle, A. Molnar and A. E. Firth for discussions. This work was supported by a Balzan Prize award and the European Research Council Advanced Investigator Grant ERC-2013-AdG 340642 TRIBE. B.Y.W.C. was supported by an EMBO long-term postdoctoral fellowship and a Sir Henry Wellcome Fellowship [096082]. D.C.B. is the Royal Society Edward Penley Abraham Research Professor.

## Author contributions

B.Y.W.C. and D.C.B. conceived and designed the research. B.Y.W.C performed and analysed the data. M.J.D., A.J.G. and J.H. performed all the LC-MS/MS sample processing and iSPY analysis. B.Y.W.C. and D.C.B. wrote the manuscript.

## Conflict of interest

The authors declared that they have no conflict of interest.

---

**Supplementary Figure 1.**
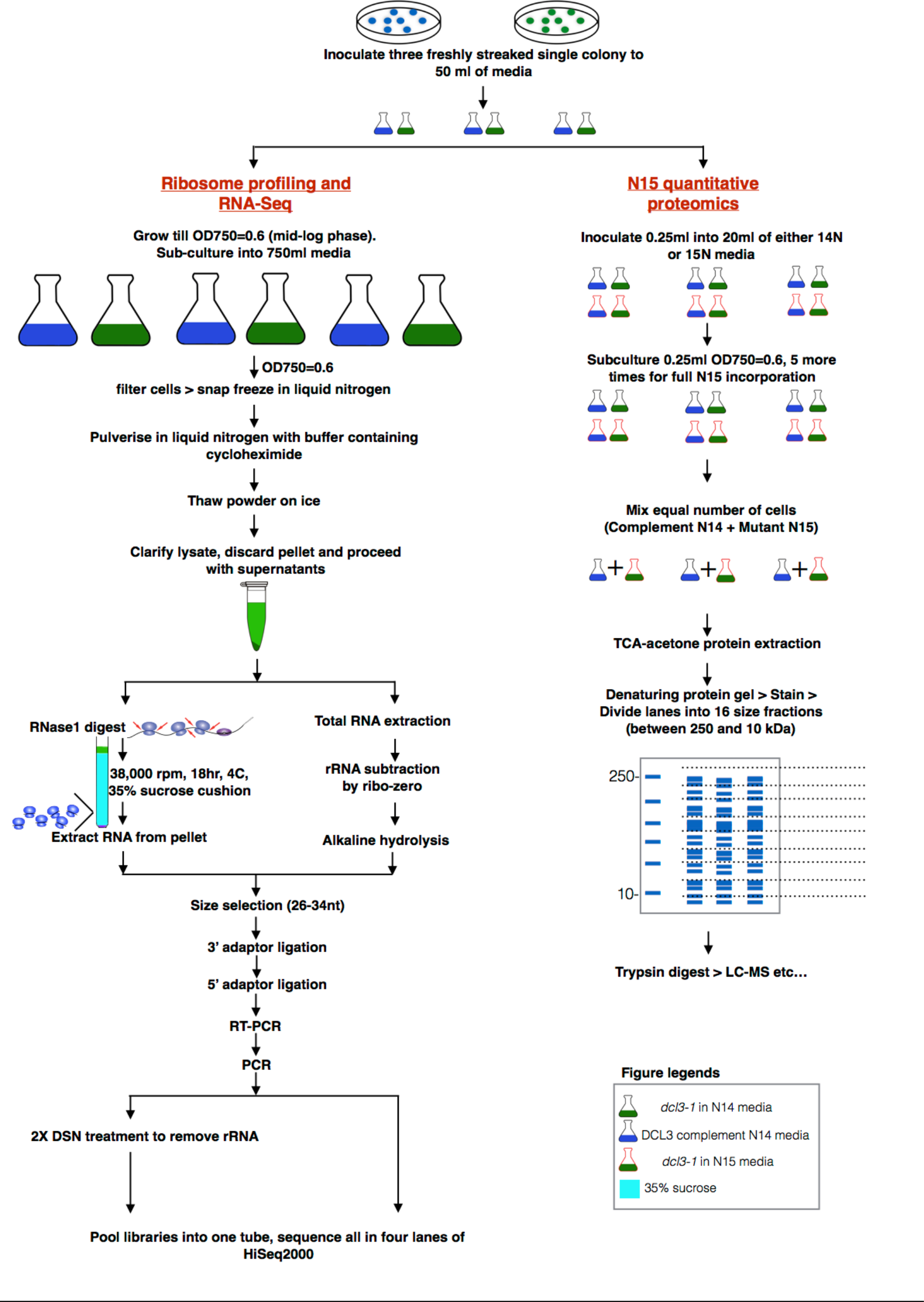
Experimental workflow. Three independent single colonies from freshly streaked *Chlamydomonas dcl3-1* (green) or complement (blue) were inoculated into 50 mL of TAP media and grown until OD750 = 0.6 (mid-log phase). 0.25 mL of each culture was used for N15 incorporation for whole cell proteomics and the remaining culture was used to sub-culture 750 mL of TAP for ribosome profiling.

**Supplementary Figure 2.**
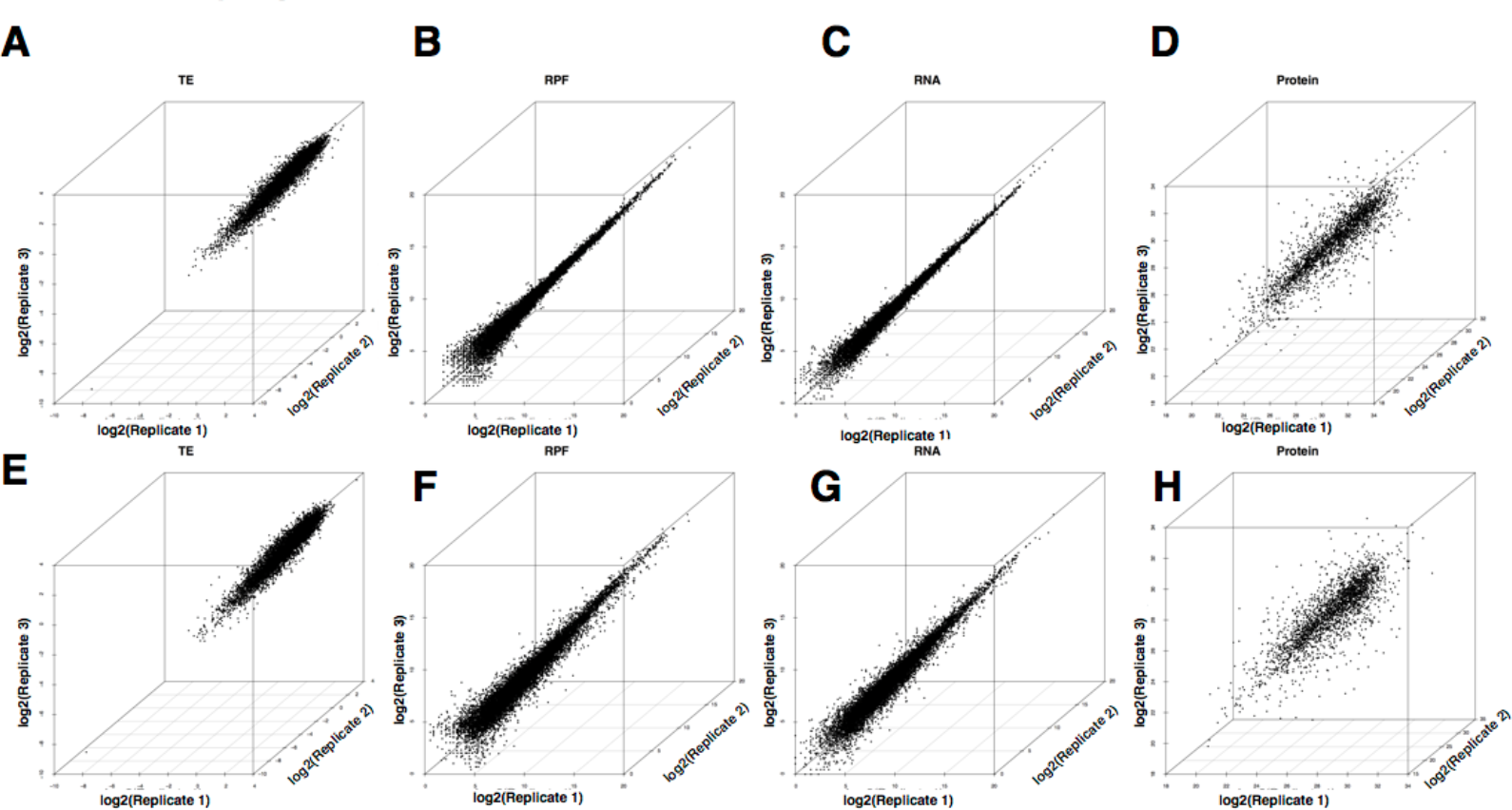
Reproducibility of TE, ribosome profiling, RNA-Seq and N15 Proteomics. (A)-(D) Correspondence between biological triplicates for DCL3-complements. (E)-(H) Correspondence between biological triplicates for *dcl3-1*.

**Supplementary Table 1:** Number of reads mapping to nuclear-encoded transcripts for each library (Phytozome 281).

|  |  | biorep 1 | biorep 2 | biorep 3 |
| --- | --- | --- | --- | --- |
| RiboSeq | Complement | 368,613 | 291,373 | 444,768 |
|  | DCL3 | 515,717 | 461,013 | 590,953 |
| RNA-seq | Complement | 908,865 | 1,114,867 | 1,223,427 |
|  | DCL3 | 1,166,393 | 1,183,071 | 679,375 |

**Supplementary Figure 3:**
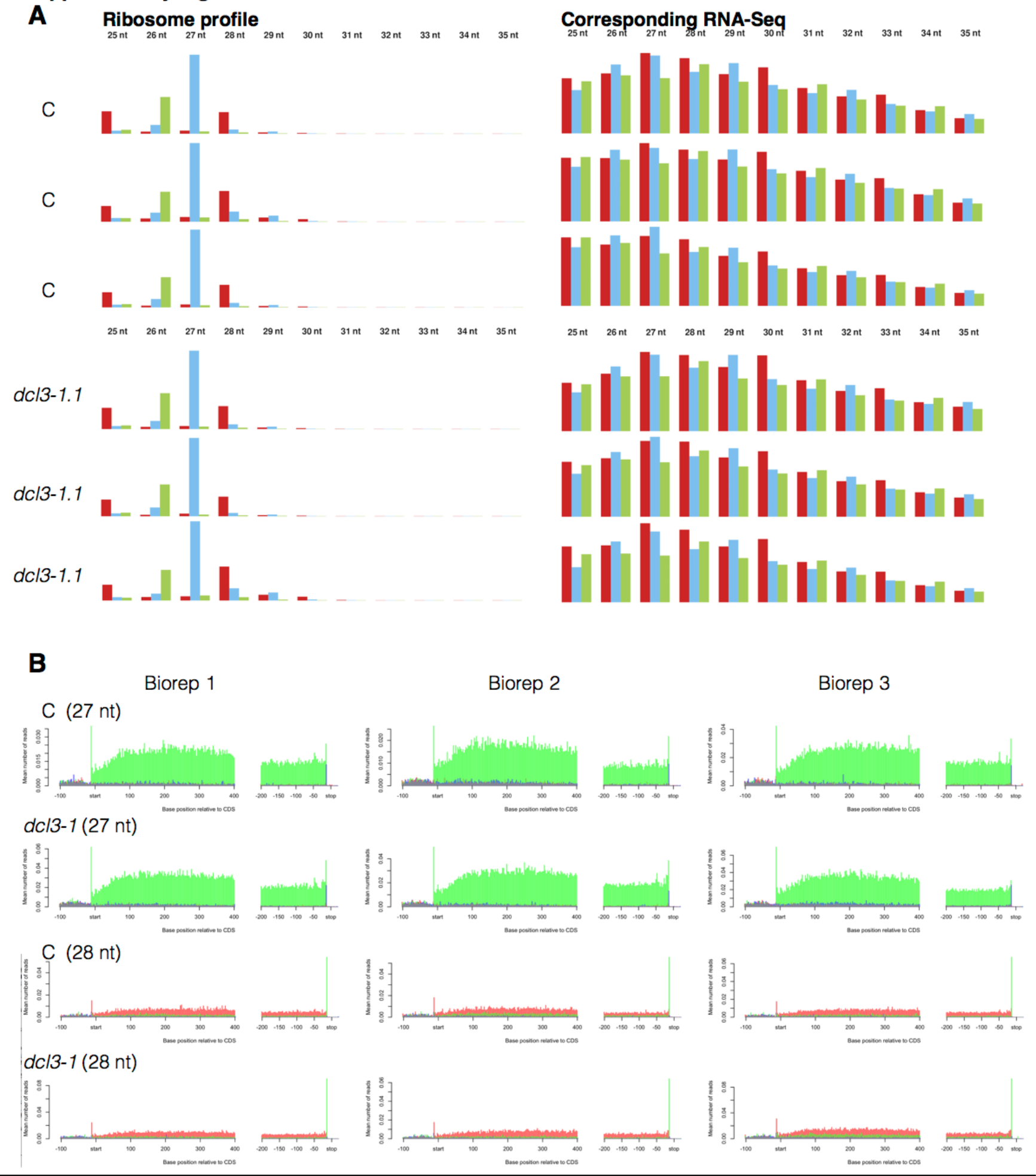
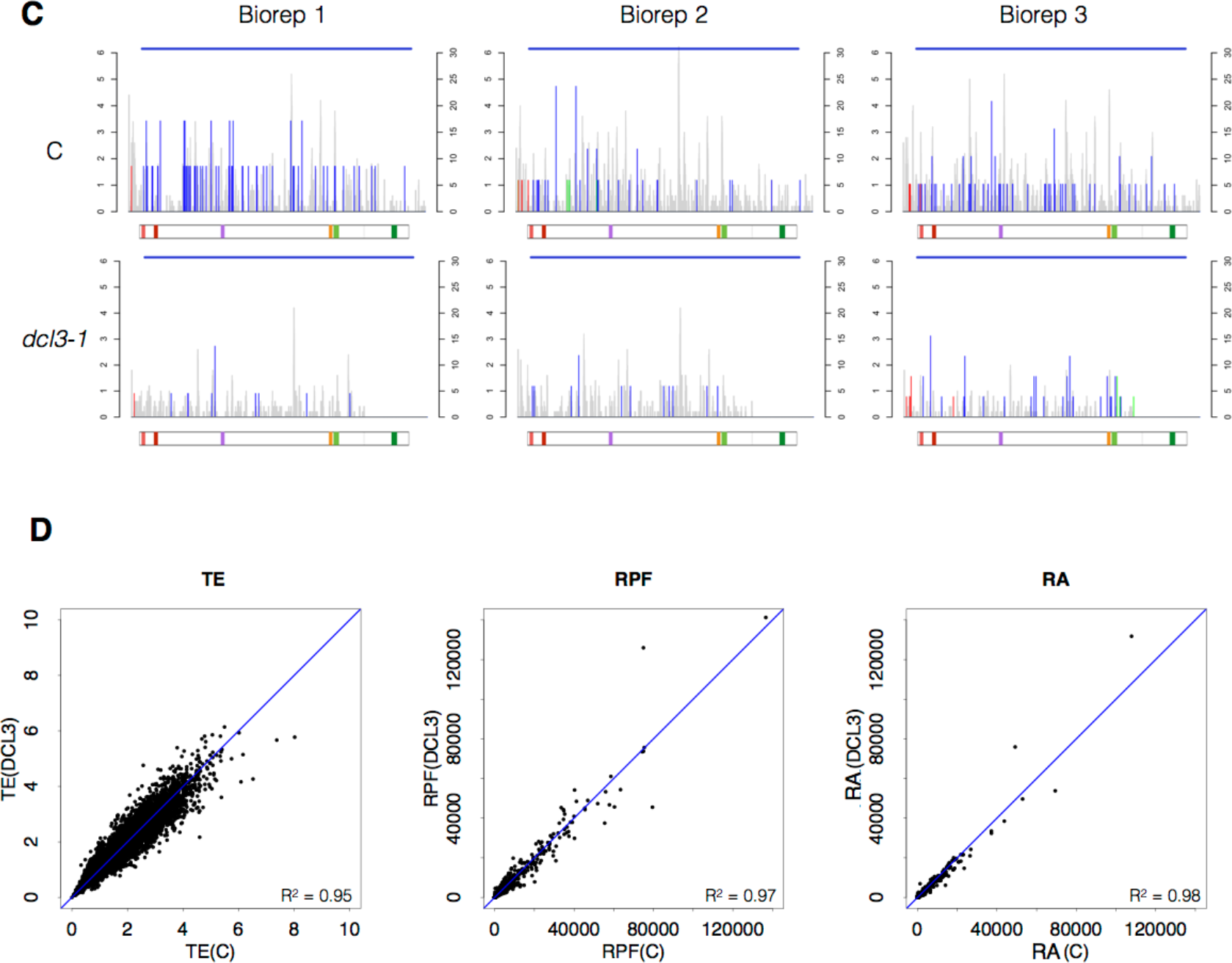
Generation of precise ribosome profiling data: (A) Histogram of positions for all biological triplicates to which the 5’ ends of ribosome profile footprints (RPFs) and corresponding RNA-Seq reads map, respectively, as a function of read size class (nt), for reads mapping to the interior region of nuclear-encoded coding ORFs. Red, green and blue bars indicate the proportion of reads that map to codon positions 0, 1 and 2 (respectively). (B) Histogram of 5’ end positions of 27 and 28-nt RPFs relative to start and stop codons for all biological triplicates. Reads were derived from the complement or *dcl3-1* (respectively) and summed over all transcripts. Phasing is indicated using the same colours as in panels A and B. Histograms of 5’ end positions of RPFs (coloured, left-axis) and RNA-Seq reads (grey, right-axis). (C) 27-nt reads mapped to DCL3 transcripts in all biological triplicates. The blue horizontal line indicates the re-annotated CDS (612-12,830 nt). The schematic below the plot shows the domain organisation of DCL3 which contains two DEAD/DEAH box helicase domains (light and dark red boxes), a Helicase C domain (purple box), a proline-rich domain (orange box) and two Ribonuclease III domains a and b (light and dark green boxes, respectively). The thin grey line and the corresponding red arrow indicates the Hygromycin insertion site (nt 10,193). (D) Correlation of TE, RPF and RNA (averaged over biological repeats) between dcl3-1 and C for all expressed genes. Blue lines represent a perfect correlation. Spearman correlation coefficients are indicated in bottom right corners.

**Supplementary Table 2:**
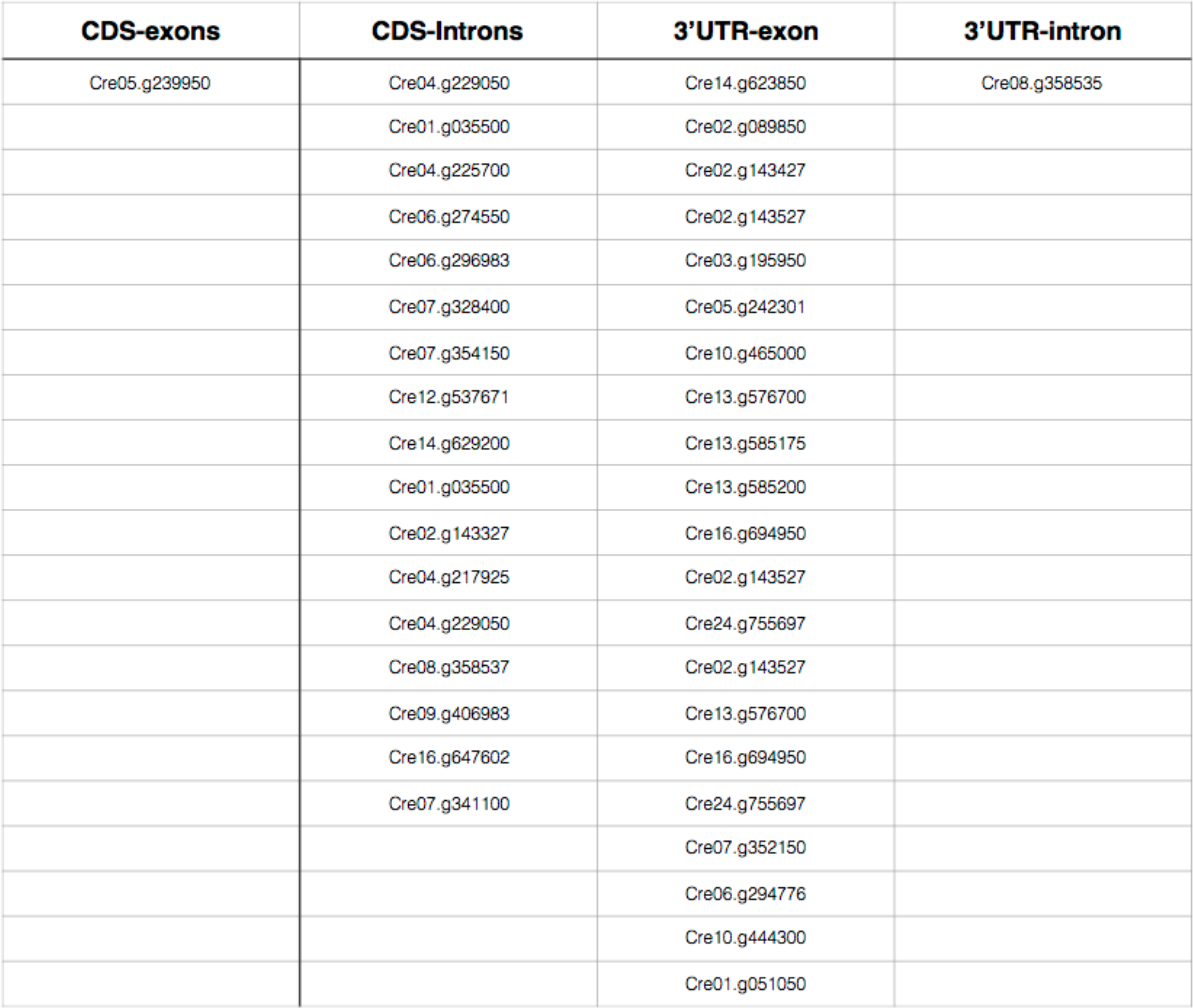
Re-annotation of miRNA precursor-containing mRNAs.

**Supplementary Figure 5:**
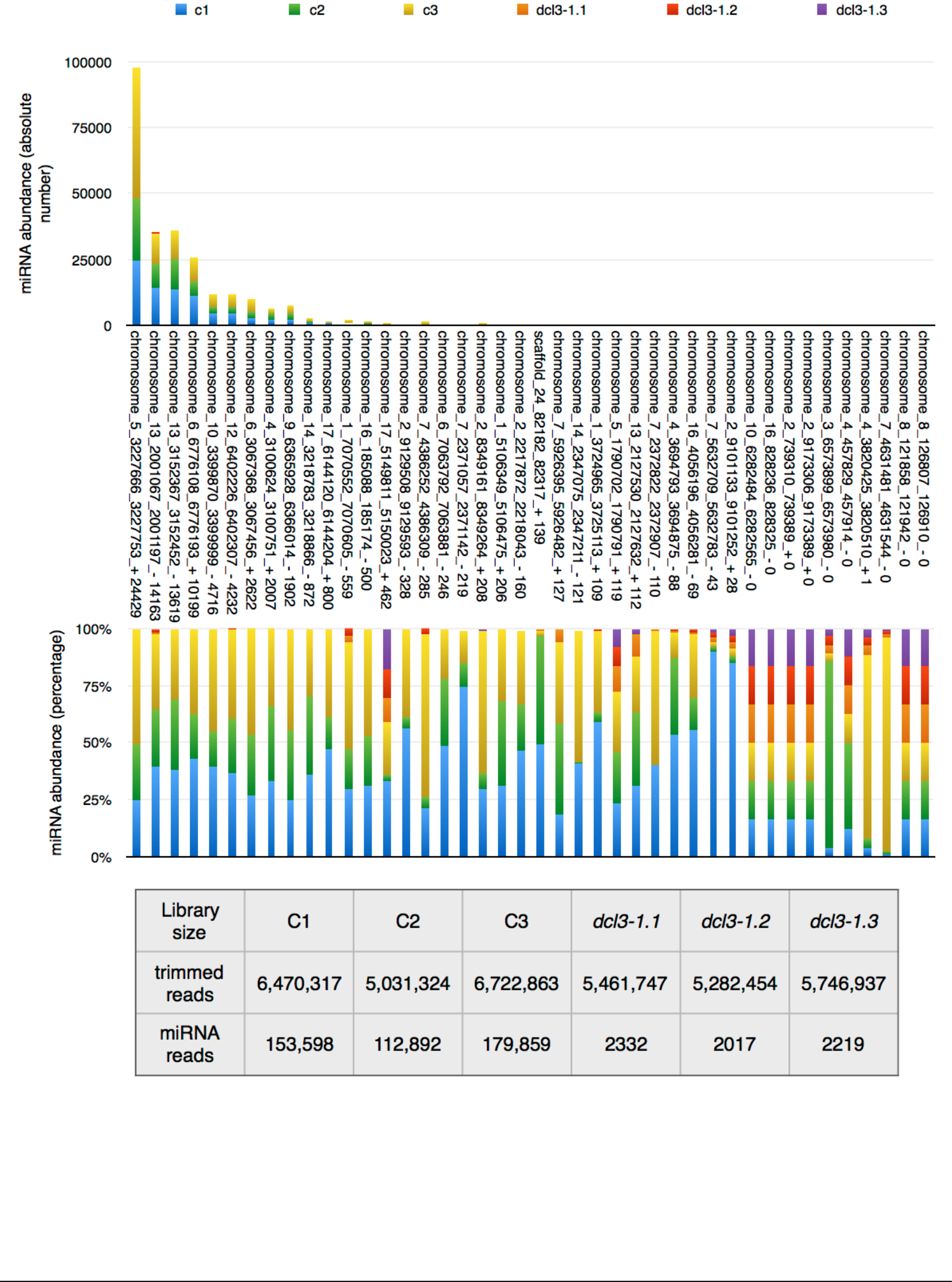
miRNA quantification. Absolute (top) and relative (bottom) quantification for all known positive-strand miRNA reads detected in all corresponding sRNA-seq libraries. Sequencing and miRNA alignment statistics for each library are in the table below.

**Supplementary Figure 6.**
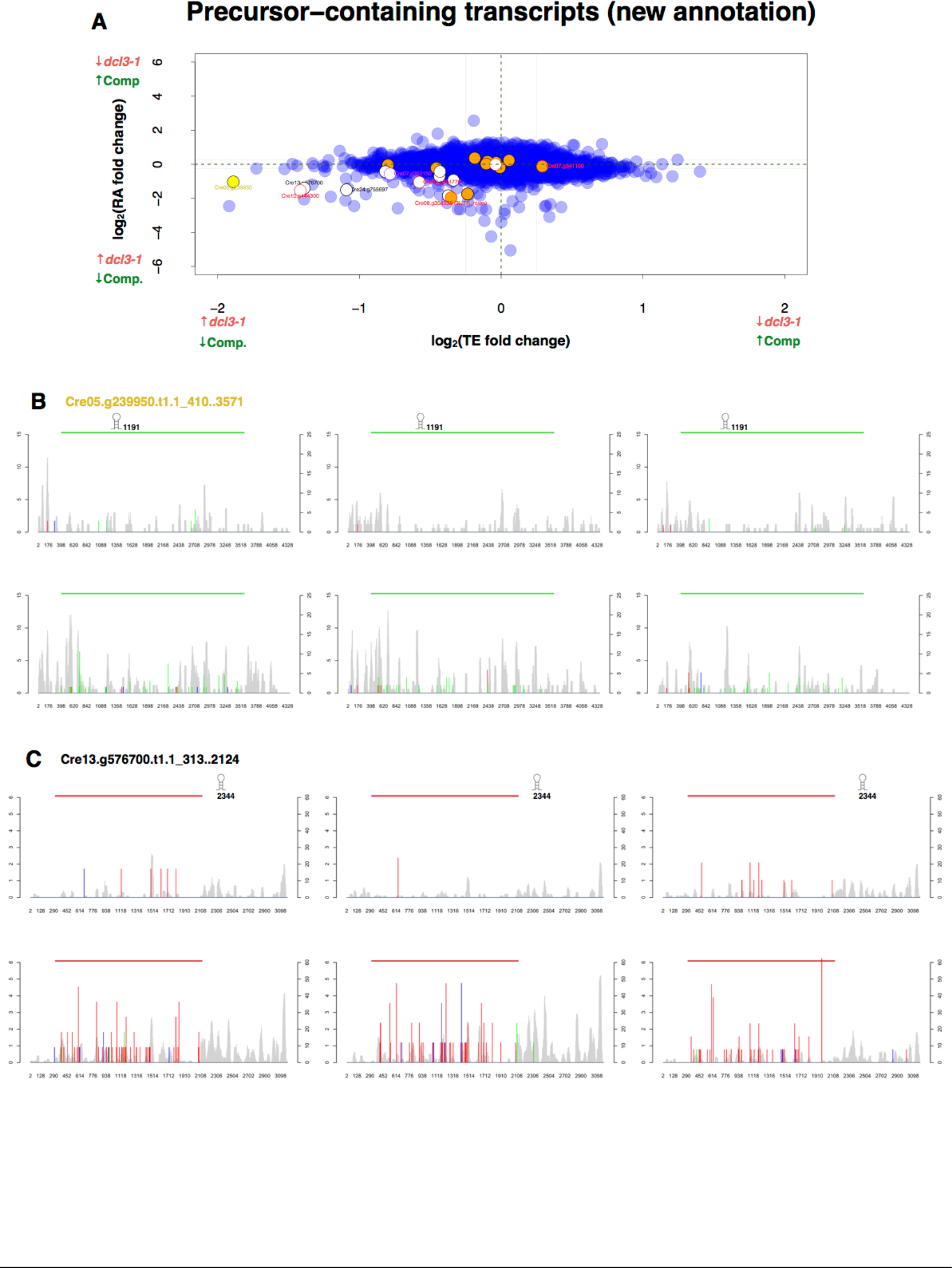
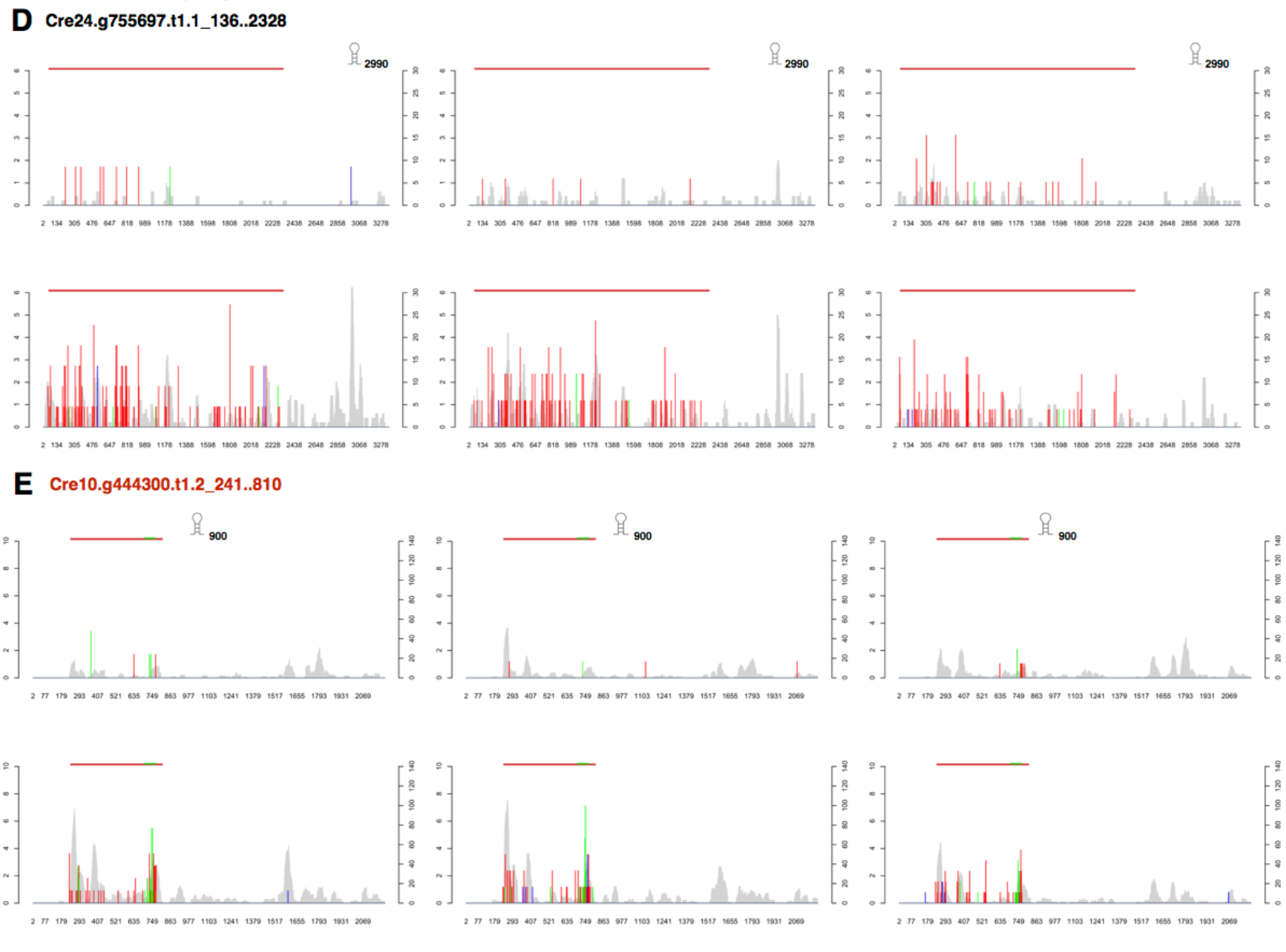
DCL3-dependent processing of miRNA occurs in the cytoplasm and down-regulates translation efficiency. (A) Scatter plot of log_2_ fold changes of all mRNAs for TE and RA fold-changes between *dcl3-1* and C. New annotation for precursor-containing transcripts: yellow circle = precursor-containing CDS, white circles = precursor-containing 3’UTRs, orange circles = precursor-containing introns, white circles with red outlines = transcripts previously annotated as non-coding transcripts but which are in fact coding and contain a miRNA precursor in the 3’UTR. Pearson correlation = 0.072 and 0.486 for intron-and exon-containing transcripts respectively. (B)-(E) Histogram of normalised 5’ end positions of 27-nt RPFs relative to start and stop codons (colour) and corresponding RNA-seq reads (grey) for miRNA-precursor containing transcripts. Reads were derived from the complement or DCL3 mutant (top and bottom in biological triplicates, respectively) and summed over all transcripts.

**Supplementary Figure 7:**
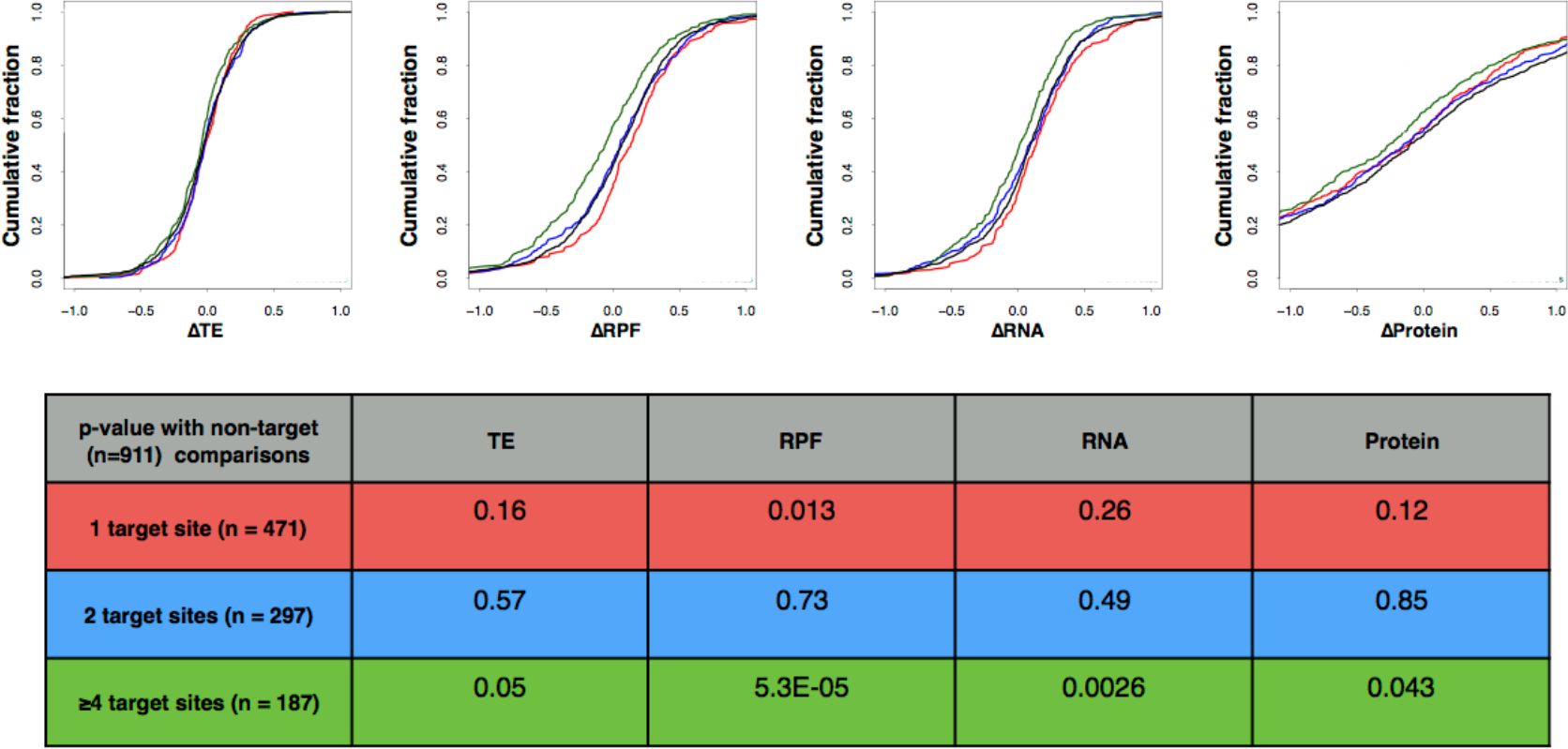

Cumulative *dcl3-1* relative to C log_2_ fold-change distributions of ΔTE, ΔRPF, ΔRA and ΔProtein for genes with both NGS and proteomic support and with 0 (black), 1 (red), 2-3 (blue) or 4 or more (green) target sites. K.S. p-values are shown in the table below.

**Supplementary table 3.**
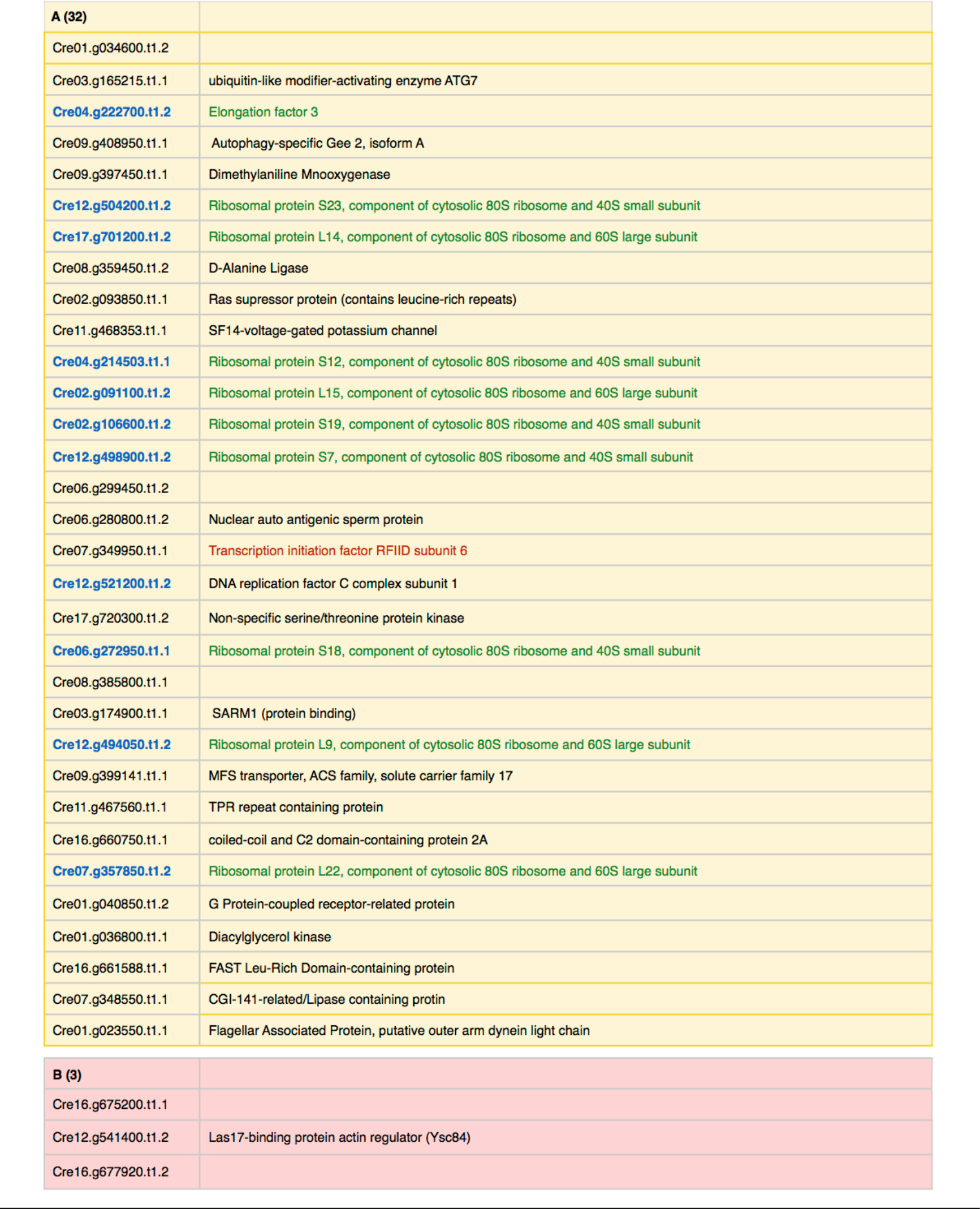
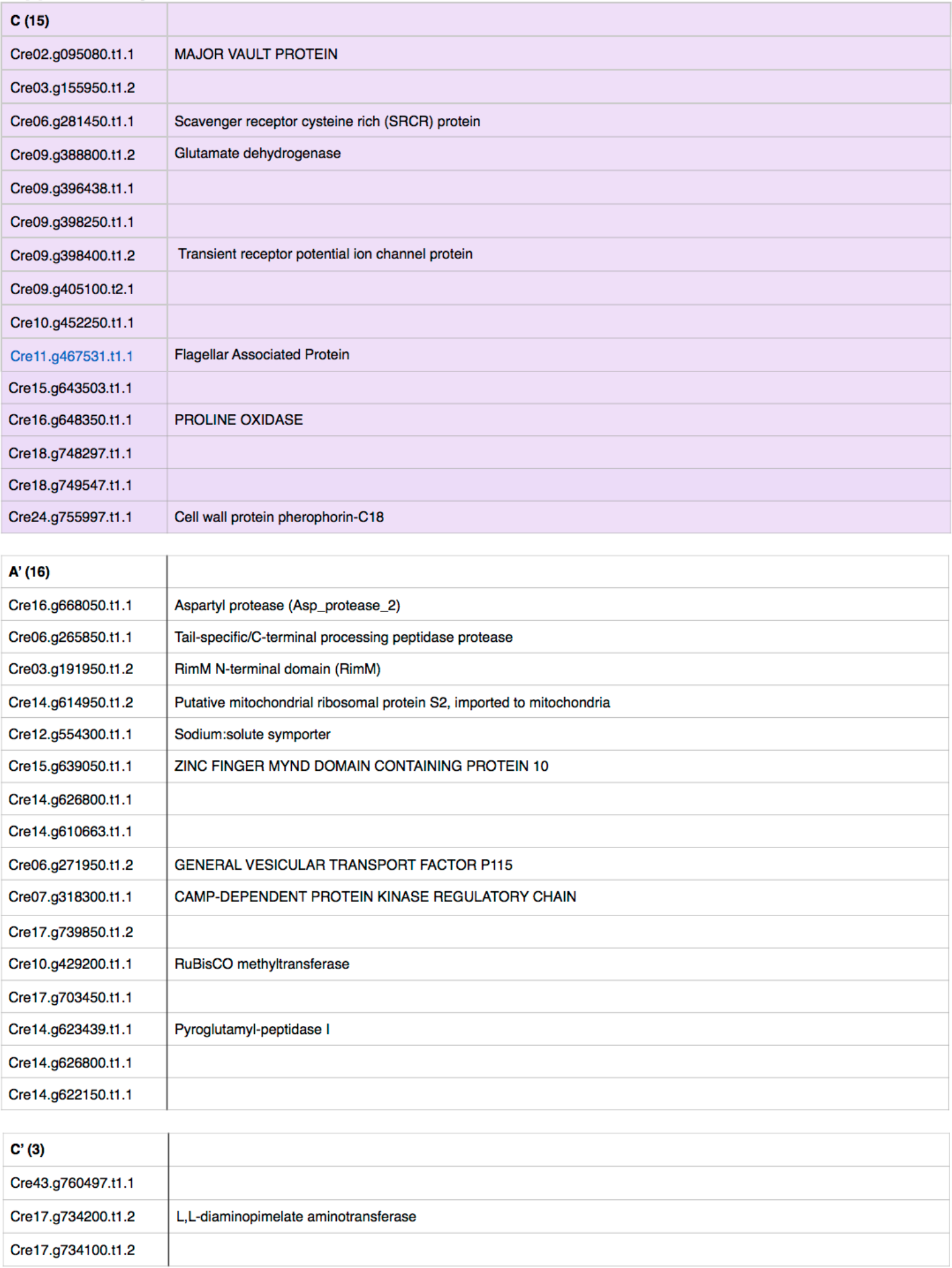
Lists of mRNAs lying within boxes A, A’, B, B’, C and C’ (Figure 4A) and their respective annotations. Annotations associated with the 80S translation machinery are highlighted in green, and other RNA binding proteins in red. Messenger RNAs with detectable protein in the N15 proteomics data are highlighted in blue

**Supplementary Figure 8.**
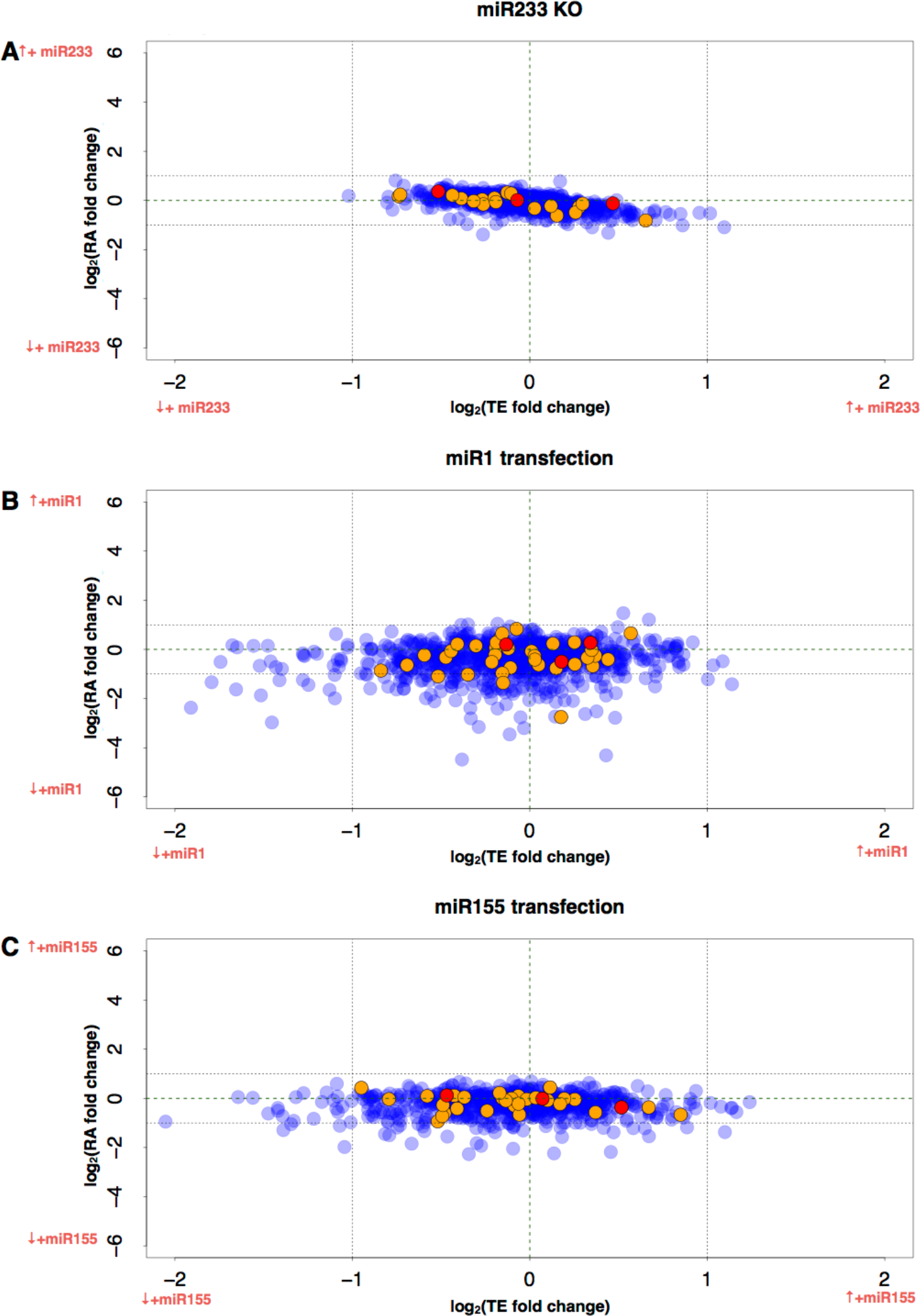

Correspondence between ΔTE and ΔRA log_2_ fold-changes after deleting miR-233 in mouse neutrophil cells (A), after introducing miR-1 to HEK293 cells (B), and after introducing miR-155 to HEK293 cells (C). Fold-change data were obtained from (Guo *et al*, 2010).

## References

Agarwal V, Bell GW, Nam JW & Bartel DP (2015) Predicting effective microRNA target sites in mammalian mRNAs. Elife 4:

Ameres SL & Zamore PD (2013) Diversifying microRNA sequence and function. Nat. Rev. Mol. Cell Biol. 14: 475–88 Available at: http://www.nature.com/nrm/journal/v14/n8/full/nrm3611.html#affil-auth

Bartel DP (2009) MicroRNAs: target recognition and regulatory functions. Cell 136: 215–33

Bazzini A a, Lee MT & Giraldez AJ (2012) Ribosome profiling shows that miR-430 reduces translation before causing mRNA decay in zebrafish. Science 336: 233– 7 Available at:

Bonnet E, He Y, Billiau K & van de Peer Y (2010) TAPIR, a web server for the prediction of plant microRNA targets, including target mimics. Bioinformatics 26: 1566–1568

Brodersen P, Sakvarelidze-Achard L, Bruun-Rasmussen M, Dunoyer P, Yamamoto YY, Sieburth L & Voinnet O (2008) Widespread translational inhibition by plant miRNAs and siRNAs. TL - 320. Science 320 VN-: 1185–1190 Media/brodersen2008.pdf%5Cnhttp://dx.doi.org/10.1126/science.1159151

Brodersen P & Voinnet O (2009) Target Recognition and Mode of Action. Nat. Rev. Mol. Cell Biol. 10: 141–148

Brosch M, Yu L, Hubbard T & Choudhary J (2009) Accurate and sensitive peptide identification with mascot percolator. J. Proteome Res. 8: 3176–3181

Chung BY, Hardcastle TJ, Jones JD, Irigoyen N, Firth AE, Baulcombe DC & Brierley I (2015) The use of duplex-specific nuclease in ribosome profiling and a user-friendly software package for Ribo-seq data analysis. Rna 21: 1731–1745

Eichhorn SW, Guo H, McGeary SE, Rodriguez-Mias RA, Shin C, Baek D, Hsu S hao, Ghoshal K, Villén J & Bartel DP (2014) MRNA Destabilization Is the dominant effect of mammalian microRNAs by the time substantial repression ensues. Mol. Cell

Gao X, Zhang F, Hu J, Cai W, Shan G, Dai D, Huang K & Wang G (2016) MicroRNAs modulate adaption to multiple abiotic stresses in Chlamydomonas reinhardtii. Sci. Rep. 6: 38228

Guo H, Ingolia NT, Weissman JS & Bartel DP (2010) Mammalian microRNAs predominantly act to decrease target mRNA levels. Nature 466: 835–40

Gutteridge A, Pir P, Castrillo JI, Charles PD, Lilley KS & Oliver SG (2010) Nutrient control of eukaryote cell growth: a systems biology study in yeast. BMC Biol. 8: 68

Hardcastle TJ & Kelly KA (2010) baySeq: Empirical Bayesian methods for identifying differential expression in sequence count data. BMC Bioinformatics 11: 422

Ingolia NT, Ghaemmaghami S, Newman JRS & Weissman JS (2009) Genome-wide analysis in vivo of translation with nucleotide resolution using ribosome profiling. Science 324: 218–23

Iwakawa H & Tomari Y (2013) Molecular Insights into microRNA-Mediated Translational Repression in Plants. Mol. Cell 52: 591–601

Korostelev A, Trakhanov S, Laurberg M & Noller HF (2006) Crystal Structure of a 70S Ribosome-tRNA Complex Reveals Functional Interactions and Rearrangements. Cell 126: 1065–1077

Lewis BP, Shih I, Jones-Rhoades MW, Bartel DP & Burge CB (2003) Prediction of mammalian microRNA targets. Cell 115: 787–98

Li S, Liu L, Zhuang X, Yu Y, Liu X, Cui X, Ji L, Pan Z, Cao X, Mo B, Zhang F, Raikhel N, Jiang L & Chen X (2013) MicroRNAs inhibit the translation of target mRNAs on the endoplasmic reticulum in arabidopsis. Cell 153: 562–574

Mallory AC, Reinhart BJ, Jones-Rhoades MW, Tang G, Zamore PD, Barton MK & Bartel DP (2004) MicroRNA control of PHABULOSA in leaf development: importance of pairing to the microRNA 5’ region. EMBO J. 23: 3356–64

Marondedze C, Groen AJ, Thomas L, Lilley KS, Gehring C, Pozo JC Del, Manzano C, Feng Y, Chen M, Manley JL, Isner JC, Nühse T, Maathuis FJ, Kwezi L, Meier S, Mungur L, Ruzvidzo O, Irving H, Gehring C, Lipp JJ, et al (2016) A Quantitative Phosphoproteome Analysis of cGMP-Dependent Cellular Responses in Arabidopsis thaliana. Mol. Plant 9: 621–623

Molnar a, Schwach F, Studholme DJ, Thuenemann EC & Baulcombe DC (2007) miRNAs control gene expression in the single-cell alga Chlamydomonas reinhardtii. Nature 447: 1126–1129

Qu X, Wen J-D, Lancaster L, Noller HF, Bustamante C & Tinoco I (2011) The ribosome uses two active mechanisms to unwind messenger RNA during translation. Nature 475: 118–121

Reis RS, Hart-smith G, Eamens AL, Wilkins MR & Waterhouse PM (2015) Gene regulation by translational inhibition is determined by Dicer partnering proteins. Nat. Plants 1: 1–6

Schirle NT, Sheu-Gruttadauria J & MacRae IJ (2014) Structural basis for microRNA targeting. Science (80-.). 346: 608–613

Schmiedel JM, Klemm SL, Zheng Y, Sahay A, Blüthgen N, Marks DS & Oudenaarden A Van (2015) Expression Noise. Science (80-.).

Siciliano V, Garzilli I, Fracassi C, Criscuolo S, Ventre S & di Bernardo D (2013) MiRNAs confer phenotypic robustness to gene networks by suppressing biological noise. Nat. Commun. 4: 2364

Valli AA, Santos BACM, Hnatova S, Bassett AR, Molnar A, Chung BY & Baulcombe DC (2016) Most microRNAs in the single-cell alga Chlamydomonas reinhardtii are produced by Dicer-like 3-mediated cleavage of introns and untranslated regions of coding RNAs. Genome Res. 26: 519–529

Yamasaki T, Voshall A, Kim EJ, Moriyama E, Cerutti H & Ohama T (2013) Complementarity to an miRNA seed region is sufficient to induce moderate repression of a target transcript in the unicellular green alga Chlamydomonas reinhardtii. Plant J. 76: 1045–1056

